# Ubiquitin-related genes are differentially expressed in isogenic lines contrasting for pericarp cell size and grain weight in hexaploid wheat

**DOI:** 10.1101/175471

**Authors:** Jemima Brinton, James Simmonds, Cristobal Uauy

## Abstract

**Background:** There is an urgent need to increase global crop production. Identifying and combining genes controlling indidviual yield components, such as grain weight, holds the potential to enhance crop yields. Transcriptomics is a powerful tool to gain insights into the complex gene regulatory networks that underlie such traits, but relies on the availability of a high-quality reference sequence and accurate gene models. Previously, we identified a grain weight QTL on wheat chromosome 5A (5A QTL) which acts during early grain development to increase grain length through cell expansion in the pericarp. In this study, we performed RNA-sequencing on near isogenic lines (NILs) segregating for the 5A QTL and used the latest gene models to identify differentially expressed (DE) genes and pathways that potentially influence pericarp cell size and grain weight in wheat.

**Results:** We sampled grains at four and eight days post anthesis and found genes associated with metabolism, biosynthesis, proteoloysis and defence response to be upregulated during this stage of grain development in both NILs. We identified a specific set of 112 transcripts DE between 5A NILs at either time point, including seven potential candidates for the causal gene underlying the 5A QTL. The 112 DE transcripts had functional annotations including non-coding RNA, transpon-associated, cell-cycle control, and ubiquitin-related processes. Many of the wheat genes identified belong to families that have been previously associated with seed/grain development in other species. However, few of these wheat genes are the direct orthologs and none have been previously characterised in wheat. Notably, we identified DE transcripts at almost all steps of the pathway associated with ubiquitin-mediated protein degradation. In the promoters of a subset of DE transcripts we identified enrichment of binding sites associated with C2H2, MYB/SANT, YABBY, AT HOOK and Trihelix transcription factor families.

**Conclusions:** In this study, we identified DE transcripts with a diverse range of predicted biological functions, reflecting the complex nature of the pathways that control early grain development. Further functional characterisation of these candidates and how they interact could provide new insights into the control of grain size in cereals, ultimately improving crop yield.

## Background

Crop production must increase to meet the demands of a global population estimated to exceed nine billion by 2050 [1]. Indeed, one in nine people currently live under food insecurity [2]. With limited opportunity for agricultural expansion, increasing yields on existing land could significantly reduce the number of people at risk of hunger [3]. It is estimated that at least a 50% increase in crop production is required by 2050 [4, 5], however current rates of yield increase are insufficient to achieve this goal [6]. It is therefore critical and urgent that we identify ways to increase crop yields.

Final crop yield is influenced by the interaction of many genetic and environmental factors. This complexity hinders its study and has meant that the mechanisms controlling this trait are not well understood. Grain weight, however, an important component of final yield, is more stably inherited and is better understood than yield itself [7]. Grain weight is mainly determined by grain size, which itself is controlled by the coordination of cell proliferation and expansion processes. Studies in both crop and model species have shown that these processes are regulated by a wide range of genes and molecular mechanisms (reviewed in [8, 9]). Control at the transcriptional level has been demonstrated, with the rice transcription factor (TF) *OsSPL16* influencing grain size through cell proliferation [10], whilst a WRKY domain TF, *TTG2,* influences cell expansion in the integument of the Arabidopsis seed [11]. Important pathways relating to protein turnover have also been identified, for example the E3 ubiquitin-ligase, *GW2,* which negatively regulates grain weight and width in rice through the control of cell division [12]. *GW2* orthologues in other species, including Arabidopsis and wheat, also act as negative regulators of seed/grain size suggesting that these mechanisms may be conserved across species [13, 14]. Other pathways/mechanisms which affect grain size include microtubule dynamics [15, 16], G-protein signalling [17, 18] and phytohormone biosynthesis and signalling [19-21].

Wheat is a crop of global importance, accounting for approximately 20 % of the calories consumed by the human population [22]. However, our understanding of the mechanisms controlling grain size remains limited in wheat, compared to rice and Arabidopsis. Comparative genomics approaches have provided some insight [13, 23] and many quantitative trait loci (QTL) associated with grain size and shape components (grain area, length and width) have been identified [24-29]. However, none of these QTL have been cloned and little is understood about the underlying mechanisms. Previously, we identified a QTL associated with increased grain weight on wheat chromosome 5A. Using BC4 near isogenic lines (NILs) we determined that the QTL acts during the early stages of grain development to increase grain length through increased cell expansion in the pericarp [29]. This and other studies [13, 30, 31] suggest that the early stages of grain/ovule development are important for determining final grain size/shape in wheat.

Transcriptomics is a powerful tool to gain insights into the complex gene regulatory networks that underlie specific traits and biological processes. Several studies have used transcriptomics approaches to look at the genes expressed during grain development in wheat [32-38]. However, these studies have mostly focused on the later stages of grain development, often focusing on starch accumulation in the endosperm. Additionally, many of these studies were performed using microarrays [33, 36, 37], which represent a fraction of the transcriptome and are unable to distinguish between homoeologous gene copies. More recent studies have used RNA-seq [35, 34], which is an open-ended platform that provides homoeolog specific resolution. However, the accuracy of RNA-seq is dependent on the availability of a high-quality reference sequence and accurate gene models. Until recently, the large (∼17 Gb) and highly repetitive nature of the hexaploid wheat genome meant that genomic resources were limited and incomplete. However, this has changed drastically in the last few years with the release of several whole genome sequences and annotations [39-42]. To date, the RNA-seq grain development studies have used either expressed sequence tags (ESTs) [35, 38] or the Chromosome Survey Sequence (CSS) [34] as references. However in hindsight, these annotations are incomplete with respect to the latest gene models [41, 39]. These novel resources provide new opportunities for more detailed and accurate transcriptomic studies in wheat.

A potential drawback of transcriptomic studies is that comparisons across varieties, tissues or time points can result in a large number of transcripts being differentially expressed. While this informs our understanding of the biological mechanisms, it is difficult to prioritise specific genes for downstream analysis. Comparative transcriptomic approaches using more precisely defined genetic material, tissues and developmental time points can aid in this by defining a smaller set of differentially regulated transcripts. For example, a comparison of the flag leaf transcriptomes of wild-type and RNAi knockdown lines of the *Grain Protein Content 1* (*GPC*) genes was used to identify downstream targets of the *GPC* TFs [43]. Similarly, the transcriptomes of NILs segregating for a major grain dormancy QTL on chromosome arm 4AL were compared and specific candidate genes underlying the QTL were identified [44]. To our knowledge, no such experiments have been performed on isogenic lines with a known difference for grain size in wheat.

In this study, we performed RNA-seq on NILs segregating for a major grain weight QTL on chromosome arm 5AL. Previously, we showed that the QTL acts during early grain development and that NILs carrying the positive 5A allele (5A+ NILs) have significantly increased thousand grain weight (TGW; 7%), grain length (4%) and pericarp cell length (10%) compared to NILs carrying the negative 5A allele (5A- NILs) [29]. The NILs carry an introgressed segment of ∼490 Mb and using recombinant inbred lines we fine-mapped the grain length effect to a 75 Mb region on the long arm of chromosome 5A according to the IWGSC RefSeq v1.0. The aim of the present study was to identify biological pathways that potentially influence grain length and pericarp cell size by using RNA-seq to identify genes that are differentially regulated between the 5A- and 5A+ NILs.

## Results

### RNA-sequencing of 5A near isogenic lines

We performed RNA-seq on whole grains from two 5A NILs which contrast for grain length [29]. We chose the time point when NILs show the first significant differences in grain length (8 days post anthesis (dpa); T2) and the preceding time point (4 dpa; T1) to capture differences in gene expression occurring during this period (Figure 1). We hypothesised that although there was no significant difference in the grain length phenotype at T1, phenotypic differences were beginning to emerge and gene expression changes influencing this may already be occurring. We obtained over 362 M reads across all 12 samples (two time points, two NILs, three biological replicates), with individual samples ranging from 15.0 M to 53.6 M reads and an average of 30.2 M reads (standard error ± 3.5 M reads) per sample (Table 1). We aligned reads to two different transcriptome sequences from the Chinese Spring reference accession, the IWGSC Chromosome Survey Sequence (CSS) [40] and TGACv1 (TGAC) [41] reference. On average across samples, 69.8 ± 0.3 % of reads aligned to the CSS reference, whilst 84.4 ± 0.2 % of reads aligned to the TGAC reference.

**Table 1.** **Mapping summary of RNA-seq samples**

| Genotype | Time point | Replicate | Reads | CSS gene models |  | TGAC gene models |  |
| --- | --- | --- | --- | --- | --- | --- | --- |
|  |  |  |  | Reads pseudoaligned | % reads pseudoaligned | Reads pseudoaligned | % reads pseudoaligned |
| 5A - | 1 | 1 | 24443658 | 17072939 | 69.85 | 20549681 | 84.07 |
| 5A - | 1 | 2 | 34441799 | 23349288 | 67.79 | 28483090 | 82.70 |
| 5A - | 1 | 3 | 23462705 | 16220597 | 69.13 | 19664859 | 83.81 |
| 5A - | 2 | 1 | 21333672 | 14839724 | 69.56 | 18052324 | 84.62 |
| 5A - | 2 | 2 | 14967302 | 10632519 | 71.04 | 12803552 | 85.54 |
| 5A - | 2 | 3 | 35522754 | 25491523 | 71.76 | 30297336 | 85.29 |
| 5A + | 1 | 1 | 19267564 | 13520181 | 70.17 | 16317352 | 84.69 |
| 5A + | 1 | 2 | 22299102 | 15479234 | 69.42 | 18780525 | 84.22 |
| 5A + | 1 | 3 | 30531539 | 20789582 | 68.09 | 25436453 | 83.31 |
| 5A + | 2 | 1 | 51637607 | 36192489 | 70.09 | 43739451 | 84.70 |
| 5A + | 2 | 2 | 53575232 | 37956887 | 70.85 | 45497914 | 84.92 |
| 5A + | 2 | 3 | 30553421 | 21604895 | 70.71 | 25984674 | 85.05 |
| Total |  |  | 362036355 | 253149858 | - | 305607211 | - |
| Mean |  |  | 30169696 | 21095822 | 69.87 | 25467268 | 84.41 |

**Figure 1:**
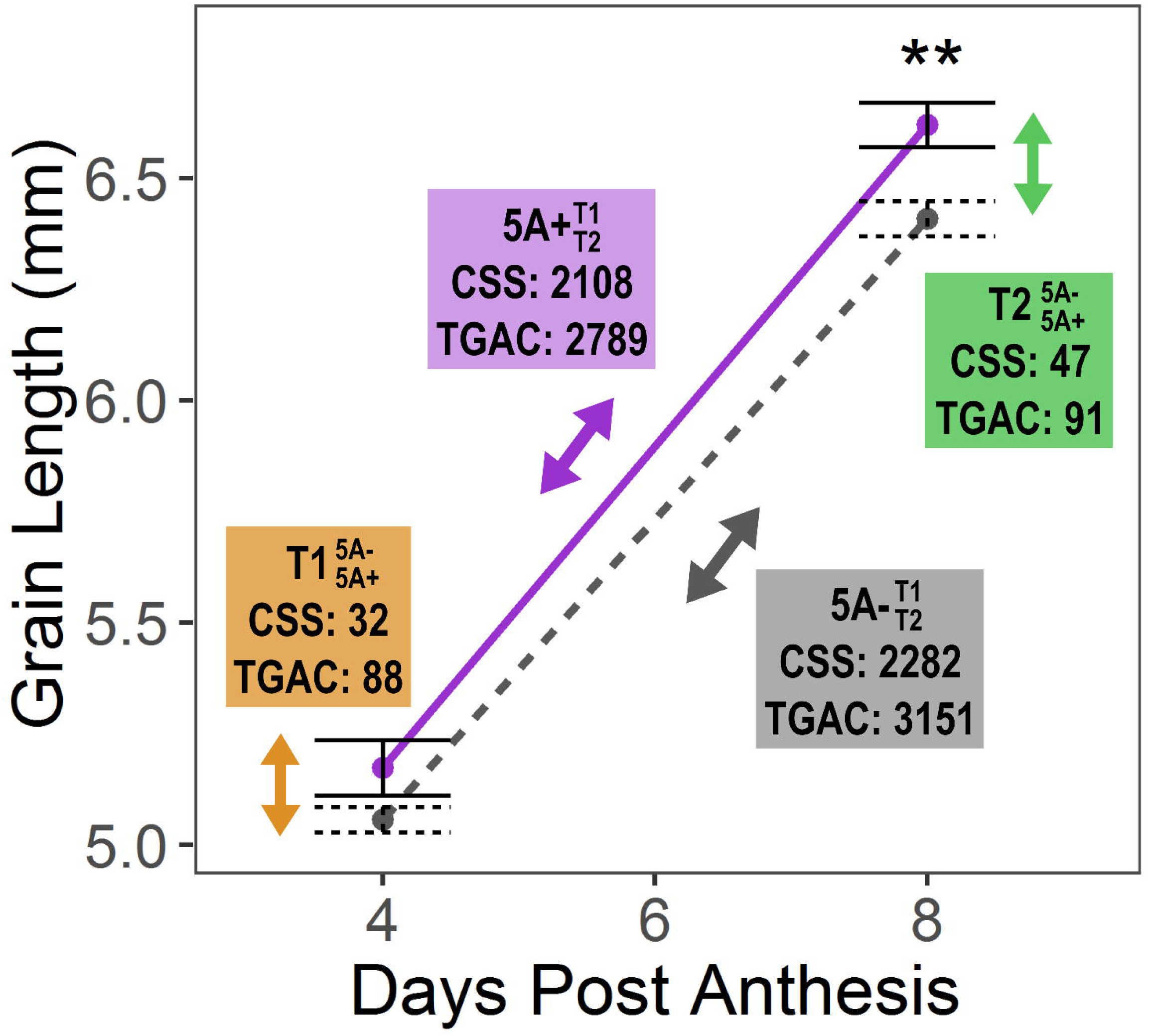
Differentially expressed genes between 5A NILs across time. RNA-seq was carried out on whole grain RNA samples taken in 4 different conditions: 5A-(short grains) and 5A+ (long grains) NILs at 4 days post anthesis (dpa; T1) and 8 dpa (T2). These were selected as the time point when the first significant difference (P < 0.01, asterisks) in grain length was observed between 5A- (grey, dashed line, short grains) and 5A+ (purple, solid line, long grains) and the preceding time point. Differentially expressed (DE) transcripts were identified for four comparisons (q-value < 0.05). Coloured boxes indicate the numbers of DE transcripts identified for each comparison using alignments to either the IWGSC Chinese Spring Survey Sequence (CSS) or the TGACv1 (TGAC) Chinese Spring reference transcriptomes. Two ‘across time’ comparisons: 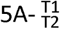 (grey box; comparing T1 and T2 samples of the 5A- NIL) and 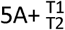 (purple box; comparing T1 and T2 samples of the 5A+ NIL), and two ‘between NIL’ comparisons: 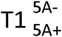 (orange box; comparing 5A- and 5A+ NILs at T1) and 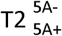 (green box; comparing 5A- and 5A+ NILs at T2).

### Comparison between Chinese Spring reference transcriptomes

We defined a transcript as expressed if it had an average abundance of > 0.5 transcripts per million (tpm) in at least one of the four conditions (2 NILs x 2 time points). This resulted in 62.5 % (64,020) and 37.1% (101,652) of the transcripts being expressed in the CSS and TGAC transcriptomes, respectively. We defined differentially expressed (DE) transcripts (q value < 0.05) using sleuth [45] and performed four pairwise comparisons: two ‘across time’ and two ‘between NIL’ comparisons. The ‘across time’ analyses consisted of a comparison between T1 and T2 samples of the 5A- NIL (hereafter symbolised as 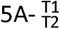; Figure 1, grey) and the corresponding comparison for the 5A+ NIL samples (hereafter 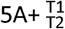; Figure 1, purple). In both cases, the T1 sample was used as the control condition, so transcripts were considered as upregulated or downregulated with respect to T1. The ‘between NIL’ analyses consisted of a comparison between the 5A-and 5A+ NILs at T1 (hereafter 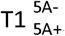; Figure 1, orange), and a comparison between the 5A- and 5A+ NILs at T2 (hereafter 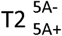; Figure 1, green). In both cases, the recurrent parent 5A- NIL was used as the control genotype. In all cases, more DE transcripts were identified in the TGAC compared with the CSS transcriptome, and similar trends were observed for both references across the four comparisons (Figure 1).

We selected the comparison with the fewest DE transcripts (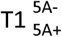; 32 and 88 DE transcripts for CSS and TGAC, respectively) to conduct a more in depth analysis of the alignments and references. For all DE transcripts from each alignment we identified the equivalent transcript/gene model in the other reference sequence using *Ensembl* plants release 35 and compared the gene models (Additional file 1). For 64 of the TGAC DE transcripts we did not identify an equivalent CSS DE transcript, either because there was no corresponding CSS gene model (47 transcripts) or the expression change between NILs was non-significant for the CSS transcript. Analogously, eleven CSS DE transcripts did not have an equivalent TGAC gene model DE, five of which were due to there being no corresponding TGAC gene model annotated. Combining both sets, we identified 42 groups of equivalent gene models, 26 of which were differentially expressed in both alignments. Comparing these 42 groups and taking into account fused and split gene models within each dataset, there were 97 gene models in both datasets (50 CSS + 47 TGAC) (Figure 2a, Additional file 1). Of these, only six were identical between the CSS and TGAC references. All other discrepant gene models fell under categories included truncations in either reference, gene models that were split/fused in one reference sequence, and gene models that differed drastically in their overall structure.

**Figure 2:**
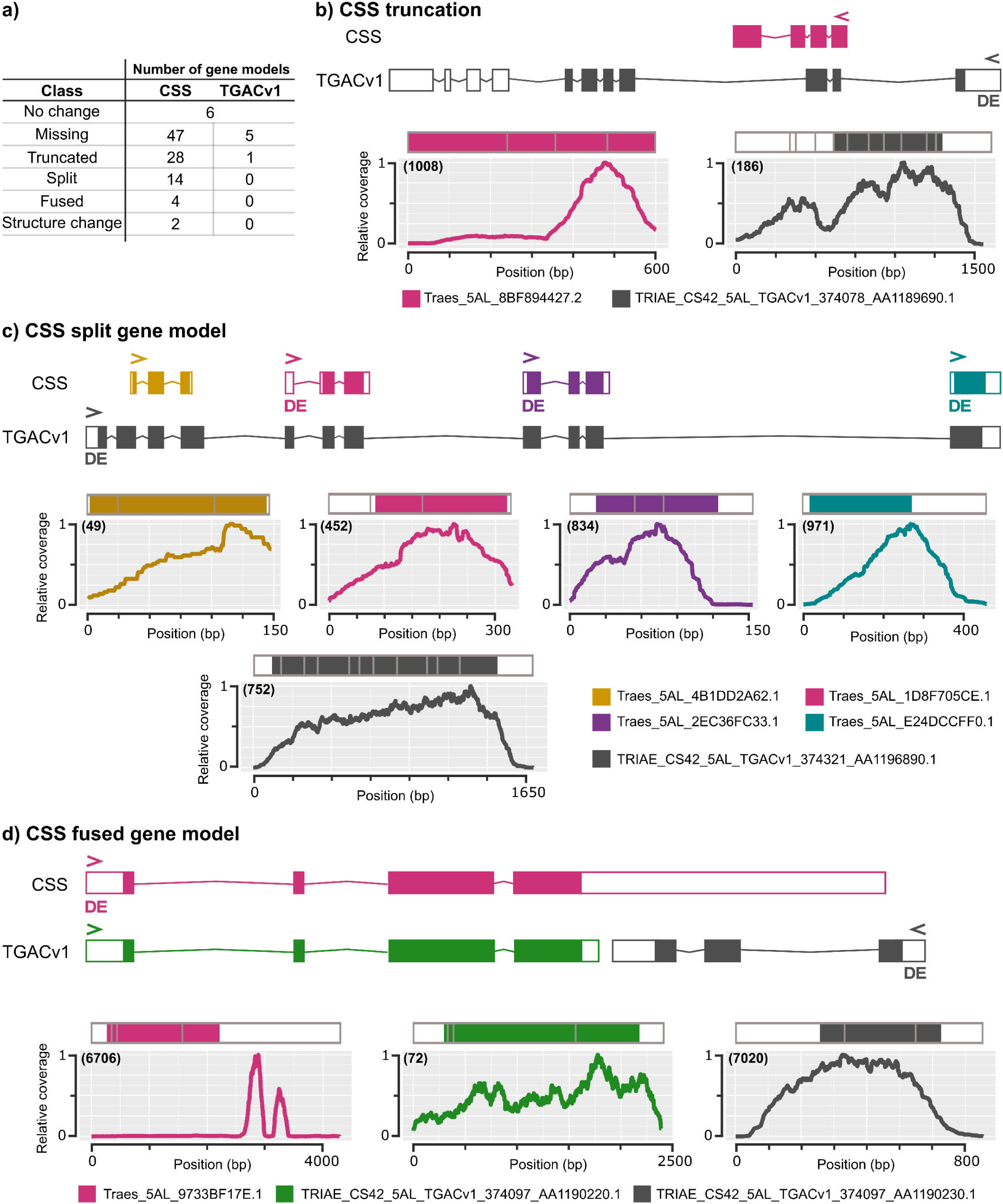
Comparison between CSS and TGACv1 gene models. a) Discrepancies identified between gene models in the CSS and TGAC reference sequences and the number of gene models falling into categories. Panels b), c) and d) show specific examples of discrepancies. In each panel, a representation of the unspliced gene model is shown with exons as coloured boxes, untranslated regions as white boxes, and introns as thin lines. Graphs show the relative read coverage across the spliced transcript with the structure represented diagrammatically directly above each graph. The number in brackets shows the maximum absolute read depth for each gene model. > and < in the gene structures indicate the direction of transcription and a ‘DE’ indicates that the gene model was differentially expressed in 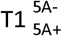 (q value < 0.05). For each panel transcript names are shown in the coloured legends.

For all discrepant gene models we used transcriptome read mapping and an interspecies comparison to determine which gene model seemed most plausible. Figure 2b shows an example of the most commonly identified discrepancy where a gene model was truncated in the CSS reference (pink) relative to the TGAC reference (grey). The DE TGAC gene model was supported by our transcriptome data as we observed read coverage across the whole gene model whilst the coverage across the CSS gene model dropped at the position where an intron is predicted in the TGAC model. Another common discrepancy was a single gene model in one reference being split into multiple gene models in the other reference. Figure 2c shows an instance where a single DE TGAC gene model comprised four separate CSS gene models. In this case, all five gene models had coverage across the entire gene body, however the single TGAC gene model was more similar to proteins from other species, suggesting that this single gene model was most likely correct. The final example (Figure 2d) shows two TGAC gene models that were fused into a single CSS gene model. The coverage across the CSS gene model was inconsistent, with most reads concentrated in the 3’ untranslated region (UTR). The two TGAC gene models had more consistent coverage across the entire gene models and were both supported by protein alignments with other species. Interestingly, only the shorter TGAC gene model was DE (Figure 2d, grey), suggesting that differential expression of the CSS gene model was driven by the reads mapping to the putative 3’ UTR rather than the coding regions of the transcript (Figure 2d, pink). Taking together the fact that a higher percentage of reads mapped to the TGAC gene models and that many more of the examined TGAC gene models were supported by interspecies comparison and expression data than the CSS gene models, we decided to continue our analysis using the alignments to the TGAC gene models only.

### Many DE transcripts during early grain development are shared between NILs

We identified 3,151 and 2,789 DE transcripts across early grain development in 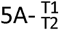 and 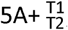, respectively (Figure 1, Figure 3a). The DE transcripts were evenly distributed across the 21 chromosomes, showing no overall bias towards any chromosome group or subgenome (Figure 3b Approximately 60% (1,832) of the DE transcripts were shared between 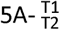 and 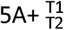 (Figure 3a) and 84% (1,532) of the shared transcripts were upregulated across time (Figure 3c). We identified 41 significantly enriched gene ontology (GO) terms in the upregulated transcripts (Additional file 2). Sixteen of the GO terms were associated with biological process and could be grouped under three parent GO terms: metabolic process (GO:0008152), defence response (GO:0006952) and biological regulation (GO:0065007) (Figure 3c). Within metabolic process we found terms associated with carbohydrate (GO:0005975) and pyruvate metabolism (GO:0006090), vitamin E (GO:0010189) and triglyceride biosynthesis (GO:0019432), mRNA catabolism (GO:0006402), proteolysis (GO:0006508) and phosphorylation (GO:0016310). Downregulated transcripts (300) were enriched for seven GO terms, four of which were associated with biological process: potassium ion transport (GO:0006813), signal transduction (GO:0007165), phosphorelay signal transduction (GO:0000160) and carbohydrate metabolism (GO:0005975). The overlap between enriched GO terms in the upregulated and downregulated transcripts (e.g. carbohydrate metabolism) suggests that different aspects of these processes are being differentially regulated during this early grain development stage.

**Figure 3:**
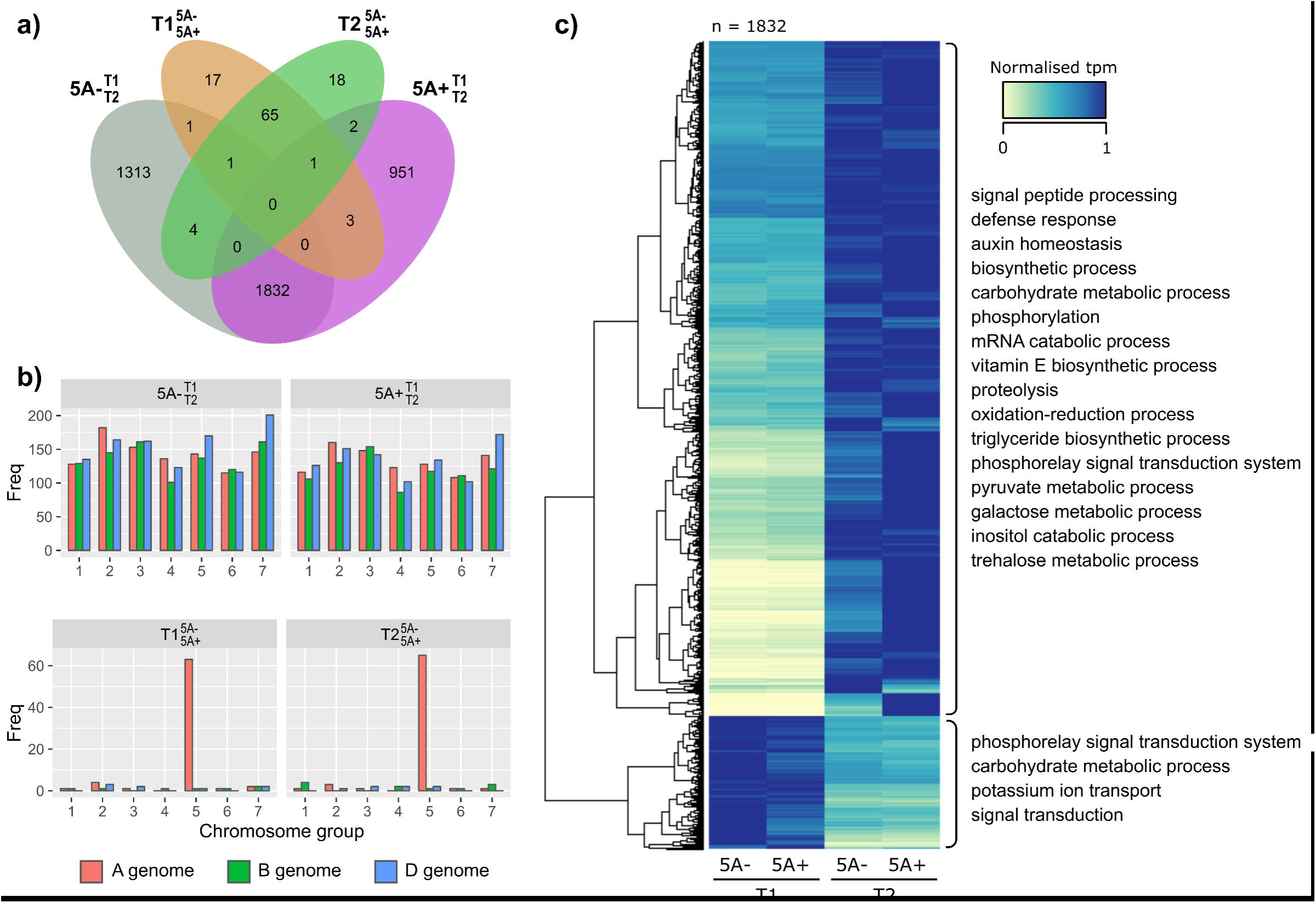
Overview of differentially expressed transcripts. a) Venn diagram of differentially expressed (DE) transcripts (q < 0.05) identified in 4 pairwise comparisons: 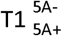 (orange), 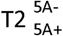 (green), 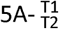 (grey) and 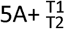 (purple). b) Number of DE transcripts located on each chromosome for all comparisons. The 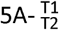 and 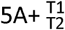 DE transcripts (top graphs) are evenly distributed across all 21 chromosomes whereas 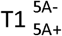 and 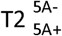 DE transcripts (bottom graphs) are concentrated on chromosome 5A. c) Heatmap of normalised tpm (transcripts per million) of common DE transcripts in 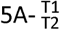 and 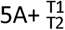 (n = 1,832). Hierarchical clustering separated these into transcripts that were upregulated (n = 1,532) and downregulated (n = 300) across time. Significantly enriched GO terms (biological function only) for each group are shown on the right of the heatmap.

We also identified many transcripts that were only DE across early grain development in one of the two genotypes (i.e. unique to either the 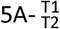 or 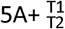 comparisons). However, many of these transcripts were borderline non-significant in the opposite genotype comparison illustrated by the fact that the distributions of q-values were skewed towards significance (Additional file 3). Additionally, the uniquely DE transcripts were enriched for GO terms similar to the shared transcripts (Additional file 2). Some GO terms, however, were only enriched in the uniquely DE transcripts, for example, cell wall organisation or biosynthesis (GO:0071554) and response to abiotic stimulus (GO:0009628). Overall, these results suggests that although there were some differences between genotypes, broadly similar biological processes were taking place in the grains of both the 5A NILs at the early stages of grain development.

### DE transcripts between NILs are concentrated on chromosome 5A

We identified 88 and 91 DE transcripts between the NILs in 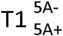 and 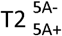, respectively, many fewer than identified in 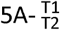 or 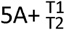. This was expected as the NILs are genetically very similar and therefore the difference in developmental stage between the T1 and T2 time points results in greater changes in gene expression. Of these 179 DE transcripts, 67 were common between 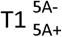 and 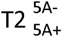, whereas 45 DE transcripts between genotypes were unique and identified only at a single time point (resulting in 112 DE transcripts between NILs at any time point). No GO terms were significantly enriched in these groups. Of the 67 common DE transcripts, 54 (80%) were located on chromosome 5A, whilst in both the T1 and T2 unique groups less than 50% were located on chromosome 5A (Figure 4a). Similar numbers of DE transcripts were more highly expressed in either genotype, with no distinct patterns observed between the unique or common groups.

**Figure 4:**
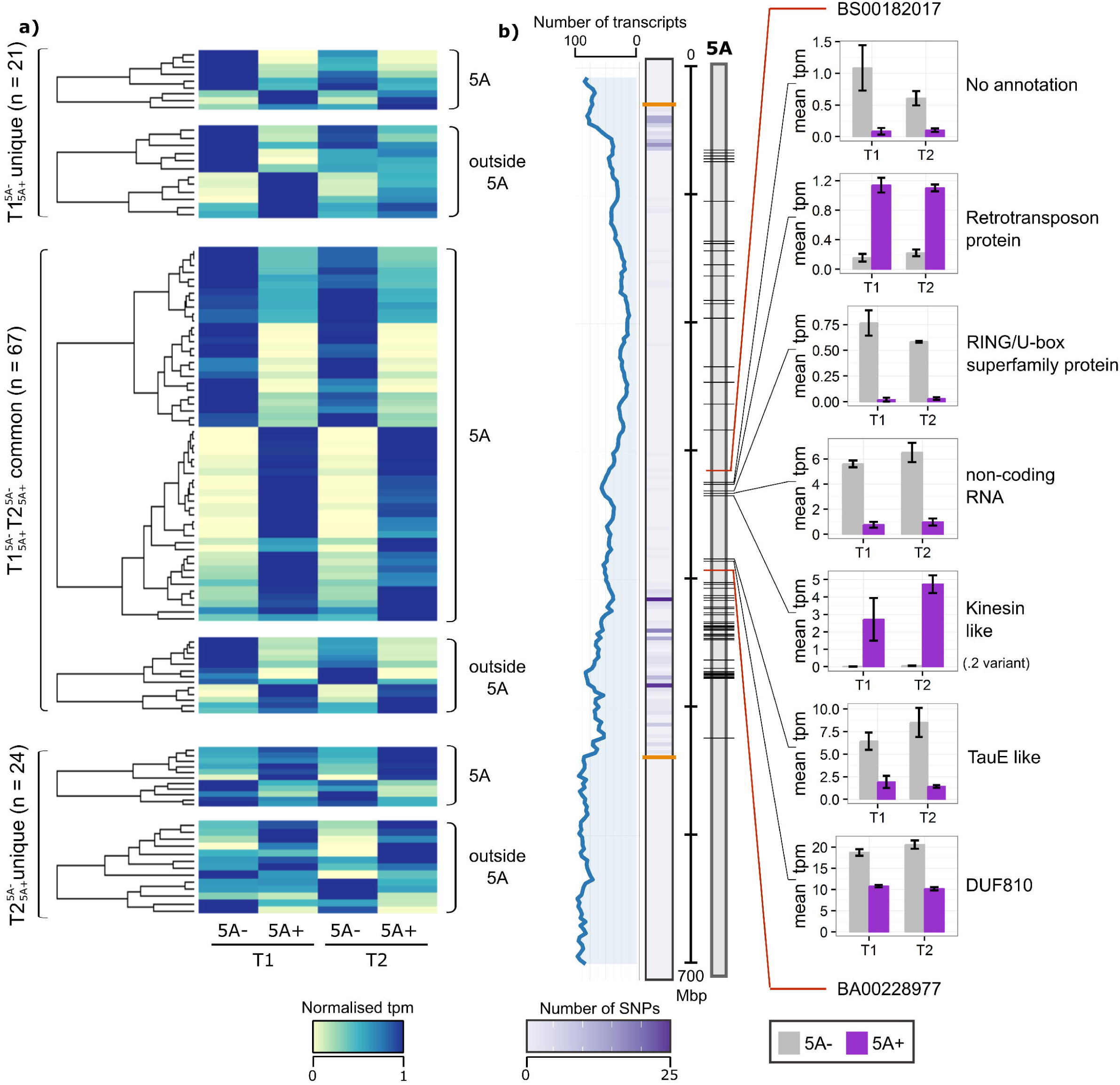
Differentially expressed transcripts between 5A NILs at T1 and T2. a) Heatmap of normalised tpm (transcripts per million) of DE (differentially expressed) transcripts between NILs (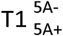 and 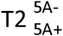 comparisons). Transcripts are first grouped based on whether they were differentially expressed at both time points (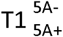 and 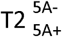 common) or at only T1 or T2 (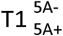 unique and 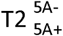 unique, respectively), and then whether they are located on chromosome 5A or not. b) Location of DE transcripts on chromosome 5A (black lines on grey rectangle). Line graph (blue) shows rolling mean of the number of transcripts located in 3 Mbp bins across chromosome 5A, alongside heatmap which shows the number of 90k iSelect SNPs between the 5A- and 5A+ NILs in 3 similar sized bins. Orange lines on the SNP heatmap define the 491 Mbp introgression which differs between then NILs. Red lines on the chromosome indicate the positions of the flanking markers of the fine-mapped region of the 5A grain length QTL (BS00182017 and BA00228977). Bar charts show the mean tpm values at T1 and T2 of DE transcripts located in the fine mapped region (5A- NILs in grey, 5A+ NILs in purple). Only one transcript variant (.2) of the kinesin-like gene is shown. Error bars are standard error of the three biological replicates.

We looked specifically at the positions of the 74 DE transcripts located on chromosome 5A and found that all were located within the 491 Mbp introgressed region of the NILs (Figure 4b). Higher numbers of DE transcripts were identified in regions of increased SNP density between the 5A NILs. Previously, we fine-mapped the grain length effect to a 75 Mbp interval on 5AL (between BS00182017 (317 Mbp) and BA00228977 (392 Mbp; [29]) and eight of the DE transcripts were located within this interval. Three of these transcripts were more highly expressed in the 5A+ NILs (5A+_high_ transcripts), two of which were transcript variants of the same gene (a kinesin-like protein; only.2 variant shown in Figure 4b). The other 5A+_high_ transcript was annotated as a putative retrotransposon protein. One of the five transcripts more highly expressed in the 5A- NIL (5A-_high_ transcript) had no annotation and the remaining four were annotated as a non-coding RNA, a RING/U-box containing protein, a TauE-like protein and a DUF810 family protein.

### DE transcripts outside of chromosome 5A are enriched in specific transcription factor binding sites

As all the DE transcripts on chromosome 5A were located within the 491 Mbp introgressed region, it is possible that the differential expression was a direct consequence of sequence variation between the NILs e.g. in the promoter regions. However, the 38 DE transcripts located outside of chromosome 5A have the same nucleotide sequence as they are identical by descent (BC_4_ NILs confirmed with 90k iSelect SNP marker data [29]). We hypothesised that these transcripts are downstream targets of DE genes, such as transcription factors (TFs), located within the 5A introgression.

To assess this, we identified transcription factor binding sites (TFBS) present in the promoter regions of these 38 DE transcripts. We identified TFBS associated with 91 distinct TF families present in this group of transcripts (Additional file 4), five of which were enriched relative to all expressed transcripts (Table 2; FDR adjusted P < 0.05). The enriched TFBS families were C2H2, Myb/SANT, AT-Hook, YABBY and MADF/Trihelix.

**Table 2:**
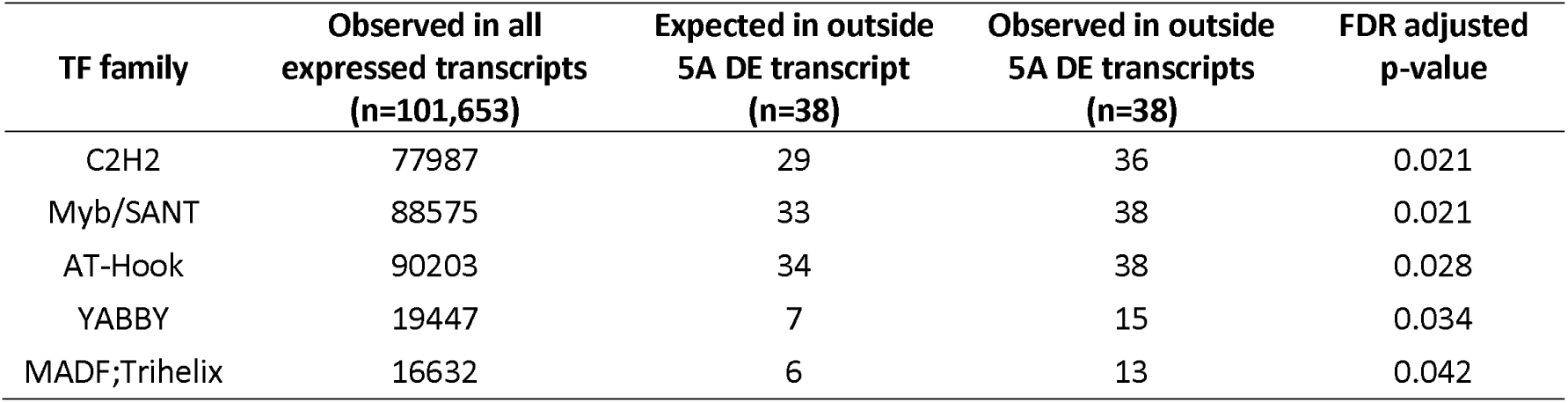
Enriched transcription factor binding sites in promoters of DE transcripts located outside of 5A. Values are the number of transcripts in which binding sites associated with the specified transcription factor (TF) family are present.

To determine potential candidates for upstream regulators we identified all TFs located within the introgressed region on chromosome 5A [46]. We identified a total of 200 annotated TFs, belonging to 35 TF families. Of these, four families corresponding to 29 TF overlapped with enriched TFBS families. Four of the 29 TFs were located within the fine-mapped grain length region on chromosome 5A, including C2H2, MYB and MYB_related TFs (Additional file 5). However, none of them were DE between NILs at the two time points.

### Functional annotation of DE transcripts

Having analysed DE transcripts between NILs based on chromosome location, we looked at the 112 DE transcripts based on their functional annotations (Additional file 6). We identified multiple categories including transcripts associated with ubiquitin-mediated protein degradation, cell cycle, metabolism, transport, transposons and non-coding RNAs (Table 3). Few categories were exclusively located on/outside 5A or had exclusively higher expression in the either the 5A- or 5A+ NIL.

**Table 3:** Categories of DE transcripts between NILs based on predicted function. Adjusted p-values displayed are based on an enrichment test of the functional categories relative to all expressed transcripts. - indicates that an enrichment test was not performed as categories were based on bespoke annotations. * includes transcripts with annotations that could not be grouped by function with other transcripts. ** only the 7 transcripts that were annotated as ubiquitin-related in the TGAC annotation were used in the enrichment test (see methods).

| Category | number of transcripts | Adjusted p-value | 5A/not 5A | NIL with higher expression: 5A-/5A+ |
| --- | --- | --- | --- | --- |
| non-coding RNA | 15 | 0.141 | 10/5 | 6/9 |
| transposon-associated | 14 | 0.008 | 4/10 | 5/9 |
| ubiquitin | 12** | 0.008 | 10/2 | 8/4 |
| cell cycle | 5 | - | 5/0 | 2/3 |
| histone-related | 5 | - | 3/2 | 3/2 |
| heat shock | 5 | - | 3/2 | 2/3 |
| protease | 4 | - | 3/1 | 3/1 |
| transport | 4 | - | 3/1 | 2/2 |
| metabolism | 5 | - | 5/0 | 4/1 |
| homeobox | 4 | 0.001 | 3/1 | 1/3 |
| cell wall | 3 | - | 2/1 | 2/1 |
| transcription | 3 | - | 2/1 | 0/3 |
| non-translating | 2 | - | 0/2 | 1/1 |
| peroxisome | 2 | - | 0/2 | 0/2 |
| other* | 20 | - | 14/6 | 11/9 |
| No annotation | 8 | - | 4/4 | 5/3 |

The category with the most DE transcripts was non-coding RNA (ncRNA, 15 transcripts), although this was not enriched relative to all expressed transcripts. All ncRNA transcripts were classed as long non-coding RNAs (>200bp, [47]) and we found that four of the ncRNAs overlapped with coding transcripts (two in the antisense direction) and one ncRNA was a putative miRNA precursor (Ta miR132-3p; [48]). We identified 13 transcripts as putative targets of Ta-miR132-3p in the TGAC reference but none of these target transcripts were differentially expressed in our dataset. The second largest transcript category was transposon-associated (14 transcripts; FDR-adjusted p = 0.008), whereas the third largest category was DE transcripts related to ubiquitin and the proteasome (12 transcripts; p = 0.008). DE transcripts annotated as homeobox were also enriched (4 transcripts; FDR-adjusted p = 0.001). Interestingly, we identified homeodomain TFBS in 27 of the 38 outside 5A DE transcripts although this was not significantly enriched (FDR-adjusted p = 0.166, Additional file 4).

The DE transcripts related to ubiquitin were of particular interest as ubiquitin-mediated protein turnover has previously been associated with the control of seed/grain size in wheat [13] and other species including rice and Arabidopsis [14, 12, 49]. The pathway acts through the sequential action of a cascade of enzymes (see Figure 5a legend) to add multiple copies of the protein ubiquitin (ub) to a substrate protein that is then targeted for degradation by the proteasome. We identified differential expression of transcripts at almost all steps of this pathway (excluding E1): two ubiquitin proteins and one ubiquitin-like protein, one E2 conjugase, six potential E3 ligase components and two putative components of the proteasome (Figure 5). In addition to these, we also identified four DE transcripts annotated as proteases (Figure 5), which are known substrates regulated by this pathway [50-52] and that influence organ size through the regulation of cell proliferation. Most of the components of the ubiquitin pathway that were differentially expressed were more highly expressed in the 5A- NIL (11/16, including proteases) (Figure 5b).

**Figure 5:**
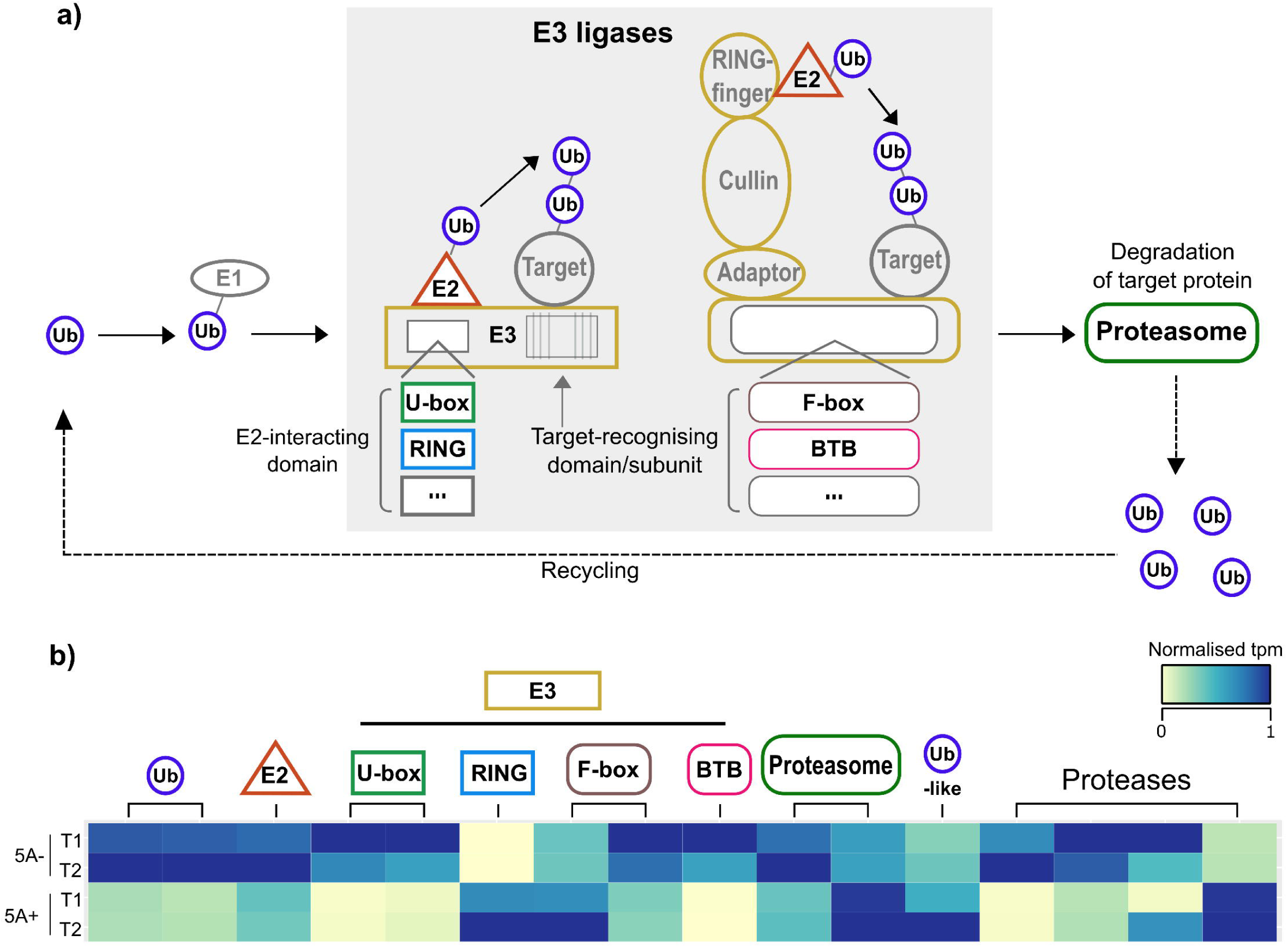
Differential regulation of the ubiquitin pathway in 5A NILs. a) Differentially expressed (DE) transcripts with functional annotations related to ubiquitin-mediated protein turnover were enriched relative to the whole genome (a). This pathway acts to add multiple copies of the protein Ubiquitin (Ub) to a substrate protein through the sequential action of a cascade of three enzymes: E1 (Ub-activating enzymes), E2 (Ub-conjugating enzymes) and E3 (Ub ligases). The tagged substrate is then targeted for degradation by the 26S proteasome and the Ub proteins are recycled. The E3 ligases are the most diverse of the three enzymes and both single subunit proteins and multi-subunit complexes exist. A subset of these classes is shown in the grey box in (a), selected based on the annotations of DE transcripts. Single subunit E3 ligases have an E2-interacting domain (e.g. U-box, RING, etc. (…)) and a substrate-recognising domain. Multi-subunit complexes also have E2-interacting complexes and substrate-recognising subunits (e.g. F-box, BTB, etc. (…)). In the context of organ size control, some proteases have been identified as downstream targets of this pathway (e.g. DA1, UBP15 [50, 51]). b) Heatmap of normalised tpm of DE transcripts associated with ubiquitin, the proteasome and proteases.

## Discussion

In this study, we performed RNA-seq on the grains of 5A NILs with a known difference in pericarp cell size, grain length and final grain weight. We previously determined that the first phenotypic differences between NILs arose during early grain development [29]. We hypothesised that differences in gene expression between NILs during these early stages would allow us to identify specific genes and pathways that that affect pericarp cell size and grain size at the transcriptional level.

### The importance of a high-quality reference sequence

We initially mapped the RNA-seq data to two different reference transcriptomes: CSS and TGAC. We found that TGAC outperformed the CSS transcriptome both in term of the number of reads that aligned and in the gene models themselves. This was most likely due to the significant improvement in terms of sequence contiguity of the TGAC reference over the CSS (N50= 88.8 vs < 10 kb, respectively), allowing more accurate prediction of gene models. Our study highlights the practical importance of this improvement as we detected 64 more DE transcripts using the TGAC reference, in most cases, due to the absence of a corresponding gene model in the CSS reference (46 transcripts). We also identified cases where incorrect gene models in the CSS reference led to misleading results. For example, in the CSS fused gene model case study (Figure 2d) a single DE transcript from the CSS reference had a large accumulation of reads mapping to the 3’ UTR. This gene was the orthologue of Arabidopsis *NPY1*, which plays a role in auxin-regulated organogenesis [53] and could therefore be related to the control of grain size. However, in the TGAC reference, in addition to the *NPY1* orthologue, an alternative gene model was annotated in place of the 3’ UTR. This alternative gene model was differentially expressed whilst the *NPY1* orthologue was expressed at a very low level and was not differentially expressed.

The improvements in scaffold size, contiguity and gene annotation open up new opportunities in wheat research. Here we used the new physical sequence to assign locations to 107 of 112 DE transcripts identified between NILs, allowing us to determine which DE transcripts were located within the QTL fine-mapped interval. Likewise, the analysis of promoter sequences enabled new hypothesis generation for this specific biological process and will also aid in the understanding of how promoter differences across genomes affects the relative transcript abundance of the different homoeologs. This exemplifies the importance of correctly annotated gene models and improved genome assemblies in gaining a more accurate view of the underlying biology.

### Differential expression analysis provides an insight into the biological processes occurring in early grain development

We sampled grains at 4 and 8 dpa to encompass the developmental stage at which the first significant difference in grain length between 5A NILs is observed. During this stage, increases in grain size are largely driven by cell expansion in the pericarp [54, 55], consistent with our previous finding that increased pericarp cell size underlies the difference in final grain length. These time points are also relatively early compared to other grain related RNA-seq studies which have focused on later grain filling processes [36, 56, 35]. The ‘across time’ comparisons (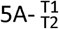 and 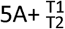) identified > 2,700 DE transcripts in each NIL, and there was a large overlap in the biological processes being differentially regulated. We found that most DE transcripts were upregulated over time and many of these were associated with metabolism and biosynthesis consistent with grains undergoing a period of rapid growth and the start of endosperm cellularisation at this stage of development [32]. Transcripts associated with proteolysis and mRNA catabolism were also upregulated across time consistent with increases in specific proteases and other hydrolytic enzymes at this stage of grain development [57]. These could be indicative of programmed cell death which occurs in both the nucellus and pericarp of the developing grain up to 12 dpa [54]. We also identified an upregulation of transcripts associated with defence response and oxidation-reduction process, consistent with previous reports of accumulation of proteins associated with defence against both pathogens and oxidative stress during the early-mid stages of grain development [58]. Transcriptional studies always have the caveat that changes in gene expression may not translate to changes in protein level [59]. However, proteomic analyses of similar stages of grain development have identified the differential regulation of similar ontologies [58, 60] suggesting that these transcriptional changes are reflective of overall protein status in the grain.

### Comparative transcriptomics as a method to identify candidate genes underlying the 5A grain length QTL

The use of highly isogenic material allowed the direct comparison of the effect of the 5A introgression on gene expression at each time point (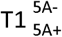 and 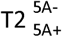). This resulted in a defined set of 112 DE transcripts between genotypes. The majority of 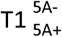 and 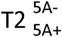 DE transcripts were located on chromosome 5A and all of these were located within the 5A introgression. This is expected given that the sequence variation in the NILs was restricted to the chromosome 5A region.

DE transcripts located within the fine-mapped interval on chromosome 5A represent good candidates for further characterisation. The kinesin-like gene and RING/U-box superfamily protein are particularly strong candidates based on their functional annotations. Previous studies have demonstrated that Kinesin-like proteins can regulate grain length and cell expansion through involvement with microtubule dynamics [15, 16, 61]. The RING/U-box protein is a putative E3 ligase, a class of enzymes which have been associated with the control of grain size (discussed in more detail later; [12, 13]).

It is premature, however, to speculate on the identity of a 5A causal gene(s) at this stage. It is difficult to predict whether DE transcripts in the fine-mapped interval are truly associated with the effect of the 5A QTL or are simply a consequence of sequence variations between the parental cultivars, i.e. ‘guilt by association’. A relevant example was the recent use of transcriptomics to define a candidate gene underlying a grain dormancy QTL (*PM19*) [44]. Subsequent studies showed that a different gene in close physical proximity (*TaMKK3*) [62] was responsible for the natural variation observed [63]. The mis-interpretation of the transcriptomics data was due to complete linkage disequilibrium between the DE *PM19* gene and the causal *TaMKK3* gene in the germplasm used in the original study. Additionally, the causal gene may not be differentially expressed between the 5A NILs and could be a result of allelic variation that alters the function of the gene independent of expression level. Ultimately, further fine-mapping of the 5A locus will be required to identify the underlying gene.

### DE transcripts outside chromosome 5A are candidates for downstream targets of the 5A QTL

We considered DE transcripts outside of chromosome 5A as candidates for downstream targets of genes located in the 5A introgression because the differential expression could not have arisen through sequence variation. These included genes located on the A, B and D genomes implying that there is cross-talk at the transcriptional level between the three genomes. We identified, in the promoters of these genes, enrichment of TF binding sites associated with TF families which have all previously been shown to play diverse roles in the control of organ development [64, 65]. For example YABBY genes, a plant specific family of TFs, play a critical role in patterning and the establishment of organ polarity [66] and fruit size [67]. Another example are the C2H2 TFs, *NUBBIN* and *JAGGED,* which are involved in determining carpel shape in Arabidopsis [68]. AT-Hook TFs play roles in floral organ development in both maize and rice [69, 70] and modulate cell elongation in the Arabidopsis hypocotyl [71]. Few of these transcription factor families have been characterised in wheat, and although these interactions need to be experimentally validated, they could be potential targets for the manipulation of grain size.

### DE transcripts have functions related to the control of seed/organ size

Studies in species such as rice and Arabidopsis have shown that seed size is regulated by a complex network of genes and diverse mechanisms, ultimately through the coordination of cell proliferation and expansion (reviewed in [8, 9]). 5A+ NILs have significantly longer pericarp cells, suggesting that the underlying gene influences cell expansion [29]. Genes that physically modify the cell wall have been shown to directly control cell expansion (reviewed in [72]) and we identified three DE transcripts that have potential roles in cell wall synthesis and remodelling. We also identified a number of DE transcripts associated with the cell cycle and the control of cell proliferation. During seed development, a number of cell cycle types in addition to the typical mitotic cycle are observed. One such alternative cycle type is endoreduplication, characterised by the replication of chromosomes in the absence of cell division, which is associated with cell enlargement (reviewed in [73]). Two of the DE transcripts were the closest wheat orthologues of Arabidopsis genes that have specific roles in organ development: a *GRF*-interacting factor (*GIF*) and *SEUSS* (*SEU*). In Arabidopsis, the GIF genes interact with the *GROWTH-REGULATING FACTOR* (*GRF*) TFs and act as transcriptional co-activators to regulate organ size through cell proliferation [74]. Conversely, *SEU* acts a transcriptional co-repressor and interacts with important regulators of development to control many processes, including floral organ development [75].

Seed development requires the coordination of processes across multiple tissues, namely the seed coat, endosperm and embryo. The development and growth of these tissues is inherently interlinked, and it has been proposed that the mechanical constraint imposed by the maternal seed coat/pericarp places an upper limit on the size of the seed/grain [30, 76, 29]. Epigenetic regulation appears to play an important role in the cross talk and coordination of these tissues [77]. The differential expression of 34 non-coding transcripts, transposons and histone-related transcripts between NILs could suggest a difference in epigenetic status associated with the control of pericarp cell size. Additional work to characterise these non-coding RNAs would be warranted to establish their role in grain development.

The ubiquitin-mediated control of seed/grain size has been documented in a number of species (reviewed in [78]), including wheat [13, 79]. DE transcripts associated with the ubiquitin pathway were significantly enriched in the 5A NILs. The pathway tags substrate proteins with multiple copies of the ubiquitin protein through the sequential action of a cascade of enzymes: E1 (Ub activating), E2 (Ub conjugases) and E3 (Ub ligases). The ubiquitinated substrate proteins are then targeted to the 26S proteasome for degradation [80]. *GW2*, a RING-type E3 ligase, negatively regulates cell division and was identified as the causal gene underlying a QTL for grain width and weight in rice [12]. The Arabidopsis orthologue, *DA2,* acts via the same mechanism to regulate seed size in Arabidopsis [14]. Another E3 ligase, *EOD1/BB* also negatively regulates seed size in Arabidopsis [49]. In general, the E3 ligase determines the specificity for the substrate proteins [80] and *DA2* and *EOD1* may have different substrate targets, however they converge and both target the ubiquitin-activated protease DA1. DA1 also negatively regulates cell proliferation and acts syngergistically with both DA2 and EOD1, although it is not clear whether the two E3 ligases act via independent genetic pathways or as part of the same mechanism [14, 81, 50]. *UBP15* (a ubiquitin specific protease) is a downstream target of this pathway and conversely acts as a positive regulator of seed size through the promotion of cell proliferation [51]. Other ubiquitin-associated regulators of organ/grain size have been identified, including components of the 26S proteasome, enzymes with deubiquitinating activity and proteins that have been shown to bind ubiquitin *in vitro* [82, 52, 83]. The DE transcripts associated with this pathway are not direct homologs of these previously characterised genes. As such the functional characterisation of these putative novel components could provide new insights into the ubiquitin-mediated control of grain size in cereals.

## Conclusions

In this study we have both generated candidates for the causal gene underlying the 5A QTL, and have also identified potential downstream pathways controlling grain size. A subset of these candidates is being tested functionally using TILLING mutants [84] and other approaches to provide novel insights into the control of grain size in cereals. Ultimately identifying the individual components and pathways that regulate grain size and understanding how they interact will allow us to more accurately manipulate final grain yields in wheat.

## Methods

### Plant material

The 5A BC_4_ NILs used in this study have been described previously [29]. Briefly, the NILs were generated from a doubled haploid population between the UK cultivars ‘Charger’ and ‘Badger’ using Charger as the recurrent parent. The NILs differ for an approximately 491 Mbp interval on chromosome 5A. We used one genotype each for the 5A- (Charger allele, short grains) and 5A+ NIL (Badger allele, long grains). Plants were grown in 1.1 x 6 m plots (experimental units) in a complete randomised block design with five replications, and spikes were tagged at full ear emergence [29]. The three blocks with the most similar flowering time were used for sampling. Plants were sampled at 4 (time point 1: T1) and 8 (time point 2: T2) days post anthesis (dpa) based on the 2014 developmental time course outlined in [29]. For each genotype, we sampled three grains from three separate spikes from different plants within the experimental unit. Each biological replicate therefore, consisted of the pooling of nine grains per genotype. Grains were sampled from the outer florets (positions F1 and F2) from the middle section of each of the three spikes. Grains were removed from the spikes in the field, immediately frozen in liquid nitrogen and stored at −80°C. In total, three biological replicates (from the three blocks in the field) were sampled for each NIL at each time point.

### RNA extraction and sequencing

For each of the three biological replicate, the nine grains were pooled and ground together under liquid nitrogen. RNA was extracted in RE buffer (0.1 M Tris pH 8.0, 5 mM EDTA pH8.0, 0.1 M NaCl, 0.5% SDS, 1% β-mercaptoethanol) with Ambion Plant RNA Isolation Aid (Thermo Fisher Scientific). The supernatant was extracted with 1:1 acidic Phenol (pH 4.3):Chloroform. RNA was precipitated at 80C by addition of Isopropanol and 3M NA Acetate (pH 5.2). The RNA pellet was washed twice in 70% Ethanol and resuspended in RNAse free water. RNA was DNAse treated and purified using RNeasy Plant Mini kit (Qiagen) according to the manufacturer’s instructions. RNA QC, library construction and sequencing were performed by the Earlham Institute, Norwich (formerly The Genome Analysis Centre). Library construction was performed on a PerkinElmer Sciclone using the TruSeq RNA protocol v2 (Illumina 15026495 Rev.F). Libraries were pooled (2 pools of 6) and sequenced on 2 lanes of a HiSeq 2500 (Illumina) in High Output mode using 100bp paired end reads and V3 chemistry. Initial quality assessment of the reads was performed using fastQC [85].

### Read alignment and differential expression analysis

Reads were aligned to two reference sequences from the same wheat accession, Chinese Spring: the Chromosome Survey Sequence (CSS; [40] downloaded from *Ensembl* plants release 29) and the TGACv1 reference sequence [41]. We performed read alignment and expression quantification using kallisto-0.42.3 [86] with default parameters, 30 bootstraps (-b 30) and the –pseudobam option. Kallisto has previously been shown to be suitable for the alignment of wheat transcriptome data in a homoeolog specific manner [87].

Differential expression analysis was performed using sleuth-0.28.0 [45] with default parameters. Transcripts with a false-discovery rate (FDR) adjusted p-value (q value) < 0.05 were considered as differentially expressed. Transcripts with a mean abundance of < 0.5 tpm in all four conditions were considered not expressed and were therefore excluded from further analyses. For each condition the mean tpm of all three biological replicates was calculated. All heatmaps display mean expression values as normalised tpm, on a scale of 0 to 1 with 1 being the highest expression value of the transcript. Read coverage for gene models was obtained using bedtools 2.24.0 genome cov [88] for each pseudobam file and then combined to get a total coverage value of each position. Coverage across a gene model was plotted as relative coverage on a scale of 0 to 1, with 1 being equivalent to the highest level of coverage for the gene model in question.

### GO term enrichment

We used the R package GOseq v1.26 [89] to test for enrichment of GO terms in specific groups of DE transcripts. We considered over-represented GO terms with a Benjamini Hochberg FDR adjusted p-value of < 0.05 to be significantly enriched.

### Functional annotation

Functional annotations of transcripts were obtained from the TGACv1 annotation [41]. Additionally, for coding transcripts we performed BLASTP against the non-redundant NCBI protein database and conserved domain database, in each case the top hit based on e-value was retained. In cases where all three annotations were in agreement, the TGAC annotation is reported. In cases where the three annotations produced differing results, all annotations are reported. Orthologues in other species such as Arabidopsis and rice were obtained from *Ensembl* plants release 36. Eight of the 112 DE transcripts had no annotation or protein sequence similarity with other species. We manually categorised the remaining 104 DE transcripts based on their predicted function. Transcripts that fell into a category of size 1 were classed as ‘other’. For the non-coding transcripts, we used BLASTN to identify potential miRNA precursors using a set of conserved and wheat specific miRNA sequences obtained from Sun et al, 2014 [48]. We used the -task blastn-short option of BLAST for short sequences and only considered hits of the full length of the miRNA sequence with no mismatches as potential precursors. We used the psRNAtarget tool (http://plantgrn.noble.org/psRNATarget/) to determine the miRNA targets.

### Identification of transcription factor binding sites

We extracted 1,000 bp of sequence upstream of the cDNA start site to search for transcription factor binding sites (TFBS). Transcripts with < 1,000 bp upstream in the reference sequence were not used in the analysis. We used the FIMO tool from the MEME suite (v 4.11.4; [90]) with a position weight matrix (PWM) obtained from plantPAN 2.0 (http://plantpan2.itps.ncku.edu.tw/; [91]). We ran FIMO with a p value threshold of <1e-4 (default), increased the max-stored-scores to 1,000,000 to account for the size of the dataset, and used a –motif-pseudo of 1e-8 as recommended by Peng et al [92] for use with PWMs. We generated a background model using the fasta-get-markov command of MEME on all extracted promoter sequences.

### Enrichment testing

To test for enrichment of different categories of transcripts relative to all expressed transcripts we performed Fisher’s exact test using R-3.2.5. For functional annotation categories, enrichment testing was only performed on categories that could be extracted using GO terms and key words based on their annotation in the TGAC reference. Only DE transcripts that could be extracted using this method were used in the enrichment tests. For example, we identified 12 DE transcripts associated with ubiquitin. The annotation of these transcripts was obtained through a combination of the TGAC annotation and manual annotation. However, only seven of these transcripts could be extracted using GO terms and key words from the whole reference annotation. Therefore, only seven transcripts were used for the enrichment test.

## Declarations

### Ethics approval and consent to participate

Not applicable

### Consent for publication

Not applicable

## Availability of data and materials

The data sets supporting the results of this article are included withing the article and its additional files. Raw sequence reads have been deposited in the NCBI Sequence Read Archive under the Bioproject PRJNA396738.

## Competing interests

The authors declare that they have no competing interests

## Authors contributions

JB designed the research, performed RNA extractions, analysed the data, performed statistical analyses and wrote the manuscript; JS coordinated the field trials and developed the germplasm used in this study; CU designed the research and wrote the manuscript. All authors read and approved the final manuscript.

## Acknowledgements

This work was funded by the UK Biotechnology and Biological Sciences Research Council (BBSRC) Grants BB/P013511/1 and BB/P016855/1 and the International Wheat Yield Partnership (grant IWYP76). JB was supported by the UK Agriculture and Horticulture Development Board (AHDB) and the John Innes Foundation. We thank David Swarbreck and Gemy Kaithakottil (Earlham Institute) for *in silico* mapping of TGACv1 gene models and to the IWGSC for pre-publication access to RefSeq v.1.0

## Additional files

Additional file 1.xlsx - **Comparison of CSS and TGAC gene models.**

Additional file 2.xlsx – **Enriched GO terms in the across time comparisons**

Additional file 3.docx – **q-value distributions of uniquely differentially expressed transcripts across time**

Additional file 4.xlsx – **Transcription factor binding sites identified in outside 5A DE transcripts**

Additional file 5.xlsx – **Transcription factors present in the 5A introgression**

Additional file 6.xlsx – **Functional annotation of DE transcripts between NILs**

